# Dynamic consensus pocket detection across molecular dynamics ensembles reveals persistent and transient druggable sites

**DOI:** 10.64898/2026.06.27.734992

**Authors:** Giulia Marigliani, Francesco Petrizzelli, Manuel Mangoni, Salvatore Daniele Bianco, Irene Orzella, Pietro Hiram Guzzi, Viviana Caputo, Tommaso Biagini, Tommaso Mazza

## Abstract

The traditional “one drug, one target” paradigm assumes that drugs interact with a single specific binding site. Modern pharmacology has proven this definition overly simplistic and, instead, recognizes that drugs operate within complex biological systems and often interact with multiple targets. In this context, proteins cannot be viewed as possessing a single functional binding site, but rather as dynamic entities capable of accommodating ligands at multiple regions, including transient and cryptic pockets. Here, we review and repurpose representative pocket detection tools across geometry-based, energy-based, and machine/deep learning approaches, originally designed to work on static conformations, to evaluate their agreement on molecular dynamics-derived conformational ensembles. Using GLUT1 protein as a dynamic transporter model and Aldose reductase as a cryptic-pocket reference system, we combine inter-tool concordance, HDBSCAN-based spatial clustering, volumetric IoU analysis, and temporal persistence scoring. Our results show that different algorithmic classes capture complementary aspects of pocket dynamics, with energy-based methods showing stronger sensitivity to transient cryptic regions and geometry-based approaches depending more strongly on pre-formed cavities. This work proposes a consensus-oriented framework for identifying conserved and transient druggable pockets in dynamic protein systems.

## INTRODUCTION

Modern drug discovery is intrinsically linked to the mechanistic understanding of protein-ligand interactions, as the precise modulation of protein function underlies most therapeutic strategies [1,2]. While rational drug design is being rapidly advanced by AI-based prediction methods and protein generative models [3,4], its success still critically depends on the accurate identification and characterization of ligand-binding pockets. These regions define where small molecules can bind and, thereby, regulate biological function [5,6]. Reliable pocket detection is therefore central to structure-based drug design, molecular docking, selectivity optimization, and the identification of alternative or allosteric binding sites [7].

Over the past three decades, a wide range of computational tools have been developed for pocket detection, broadly spanning five main methodological categories: geometry-based, energy-based, machine learning based (including deep learning), template-based, and consensus approaches [7,8]. Thus, their algorithms rely on distinct detection principles, ranging from purely structural descriptors to interaction-based scoring and data-driven learning strategies, or a consensus between them, representing an essential component of structure-based drug discovery pipelines [9]. Despite their widespread adoption, these methods have different assumptions and detection paradigms and often result in variability and limited concordance in predicted binding sites [7,8], a limitation that becomes particularly relevant in the context of protein flexibility, where distinct algorithmic strategies may emphasize different aspects of the conformational landscape. The absence of standardized benchmarks further complicates the objective selection of tools capable of providing reliable predictions within dynamic ensembles.

A common limitation of most existing pocket detection tools is that they are designed to operate on a single, static conformation of the protein [9,10]. Such representations neglect the intrinsic conformational flexibility of proteins, which modulates ligand accessibility, binding affinity, and specificity. In particular, binding sites can exhibit conformational changes ranging from subtle side-chain rearrangements to the emergence or disappearance of subpockets, or even the formation of new cavities. Molecular Dynamics (MD) simulation, which represents one of the most powerful approaches to simulate the dynamics of molecular systems over time [9–13], provides a natural framework to capture such conformational variability, enabling the characterization of pocket formation, persistence and evolution. Moreover, this dynamics perspective [77] allows the identification of cryptic pockets i.e. binding sites that are absent or only partially formed in the *apo* crystal structure but become accessible through conformational rearrangements coupled to thermal fluctuations or ligand binding. These sites typically correspond to low-population conformational states that are difficult to capture experimentally but can be sampled within MD ensembles. Since they may emerge in surface-exposed or allosterically relevant regions, distinct from the orthosteric site, cryptic pockets represent an opportunity to modulate proteins that lack classically druggable cavities. Targeting these sites with allosteric modulators has already demonstrated significant pharmaceutical value, enabling the modulation of proteins once considered undruggable [13,14]. Despite pocket detection tools being effectively able to detect and analyze these transient pockets, the choice of the drug discovery tool remains today more of an empirical approximation than a rigorous scientific selection [15].

In light of this methodological diversity, our study does not aim to provide a comprehensive benchmark of individual tools, as such comparisons are already available in the literature [8,12,16], but rather to investigate how heterogeneous pocket detection methods can be systematically integrated within a dynamic framework to characterize both stable and transient binding sites, including cryptic pockets. Adopting a philosophy similar to that of Castellana et al. [17], we focus not on the performance of individual algorithms, but on evaluating the degree of agreement across divergent detection principles when applied to dynamic conformational ensembles. Specifically, we systematically apply representative established pocket detection tools spanning geometry-based, energy-based, and machine/deep learning paradigms to analyze a library of snapshots extracted from MD trajectories of two structurally distinct proteins: GLUT1, a dynamic membrane transporter, and Aldose Reductase, a well-characterized cryptic pocket reference system. To integrate the heterogeneous outputs of these tools, we apply a dual consensus strategy combining HDBSCAN-based spatial clustering and IoU-based volumetric overlap analysis, complemented by a temporal persistence framework. This ensemble-based strategy allows us to distinguish stable, recurrently accessible pockets from transient binding regions, including cryptic sites that emerge only during specific conformational states.

## MATERIALS AND METHODS

### Pocket detection algorithms

Given the large number of available pocket detection methods, it is neither feasible nor necessary to evaluate all of them. Instead, a diverse subset of methodologies is required to provide a representative overview of current strategies for binding site prediction (**Table 1**). Existing methods can be broadly grouped into several methodological families. Geometry-based approaches, such as Fpocket [18], SiteFerret [19], EPOS [20] and KVFinder-web [21], rely on structural features such as surface concavities or solvent-excluded surfaces; KVFinder-web further provides high voxel-by-voxel precision through a dual-probe system that supports both global scans and target box definitions. Energy-based tools, including AutoSite [22], SiteHound [23,25], EasyMIFs [23], and FTSite [24], detect favorable interaction regions using virtual probe mapping. Machine learning–based approaches, including deep learning methods, such as P2Rank [26], DeepSurf [27], and DeepSite [28], use trained models to predict ligand-binding sites, while more recent deep learning tools such as PUResNetV2.0 [29] adopt a U-Net architecture for voxel segmentation. Finally, template-based methods (e.g., TM-SITE [30]) infer binding sites by structural homology to known templates, whereas consensus methods, such as MetaPocket [31], COACH [30], aggregate predictions from multiple complementary algorithms to improve prediction robustness.

**Table 1.** List of most representative pocket detection tools currently available to predict protein cavities and ligand binding sites. Abbreviations used: ML= machine learning; DL = Deep Learning.

| Tool | Criteria | Web-Server only | Open-Source | Deprecated | URL |
| --- | --- | --- | --- | --- | --- |
| Fpocket | Geometric |  | ✓ |  | https://github.com/Discngine/fpocket |
| Epos | Geometric |  | ✓ |  | https://www-cbi.cs.uni-saarland.de/software/epos_bp-ensemble-of-pockets-on-protein-surfaces-with-ballpass/ |
| SiteFerret | Geometric |  | ✓ |  | https://github.com/concept-lab/SiteFerret |
| KVFinder | Geometric | ✓ |  |  | https://github.com/LBC-LNBio/KVFinder-web-service |
| AutoSite | Energetic |  | ✓ |  | https://ccsb.scripps.edu/autosite/downloads/ |
| SiteHound | Energetic |  | ✓ |  | https://github.com/thecodingdoc/sitehound |
| EasyMIFs | Energetic |  | ✓ |  | https://github.com/thecodingdoc/easymifs |
| FTsite | Energetic | ✓ |  |  | https://ftsite.bu.edu/ |
| COACH | Template-based | ✓ |  |  | https://zhanggroup.org/COACH/ |
| ConSeq | Consensus | ✓ |  | ✓ | http://conseq.bioinfo.tau.ac.il/ |
| ConCavity | Consensus | ✓ |  |  | https://compbio.cs.princeton.edu/concavity/ |
| MetaPocket | Consensus | ✓ |  |  | https://metapocket.eml.org/ |
| P2Rank | ML |  | ✓ |  | https://github.com/rdk/p2rank |
| DeepSite | DL | ✓ |  |  | https://open.playmolecule.org/ |
| DeepPocket | DL |  | ✓ |  | https://github.com/devalab/DeepPocket |
| DeepSurf | DL |  | ✓ | ✓ | https://github.com/stemylonas/DeepSurf |
| PUResNetV2.0 | DL |  | ✓ |  | https://github.com/jivankandel/PUResNetV2.0 |

For the present comparative study, we focused on de novo pocket detection methods suitable for high-throughput analysis of MD trajectories. This choice reflects a key conceptual distinction between algorithms designed for de novo binding site discovery and those intended for the characterization of re-defined pockets [33]. Methods such as TRAPP [34–36], Epoch [37] and POVME [38–40] are highly effective for analyzing the dynamic behavior of known binding sites; however, their reliance on prior knowledge of binding site location makes them unsuitable for proteins whose druggable regions have not yet been characterized. In contrast, our objective was to identify candidate binding regions across diverse conformational ensembles without imposing prior spatial bias.

Tool selection was therefore guided by four main criteria. First, methodological diversity was required to ensure that conclusions reflected general performance trends rather than the peculiarities of a single algorithmic family. Second, tools had to be actively maintained and documented, ensuring that installation and execution were tractable within a reproducible workflow. Third, open-source local executability was required, since web-based or proprietary servers introduce rate limits, submission queues, restrictions on file size, limited control over individual frames, and reduced batch-processing capabilities, all of which are incompatible with large-scale MD analysis. Finally, selected tools needed to support high-throughput frame-by-frame trajectory analysis through command-line invocation, machine-readable output, and per-structure runtimes compatible with hundreds to thousands of snapshots. Several older discovery-oriented tools were therefore not considered further because they are difficult to deploy, often relying on outdated or deprecated software dependencies, which reduces reproducibility and hinders integration into modern computational pipelines.

Based on these criteria, six tools were selected, spanning the three principal methodological paradigms most relevant to our analysis: Fpocket and SiteFerret as geometry-based methods, AutoSite and the SiteHound as energy-based methods, and P2Rank and DeepPocket [32] as machine/deep learning-based methods. SiteHound was employed in conjunction with EasyMIFs, since SiteHound processes the molecular interaction fields computed by EasyMIFs as its primary input, forming a sequential two-step pipeline as described in the original documentation. Within each methodological category, preference was given to tools with an established presence in the benchmarking literature, as in the case of Fpocket, P2Rank, and AutoSite, while SiteFerret, DeepPocket, and SiteHound/EasyMIFs were included to broaden methodological coverage and represent less conventional but complementary approaches. For the machine/deep learning category, P2Rank and DeepPocket were specifically chosen to represent classical machine learning, based on Random Forest models, and deep learning, based on 3D-CNN architectures, respectively. Although DeepPocket builds on Fpocket as an upstream candidate generator, its CNN-based rescoring and segmentation produce a substantially reinterpreted pocket representation, as further discussed in the Discussion.

Two established tools for dynamic pocket analysis, MDpocket [45] and TRAPP [35,36], were not included in the comparative benchmark because they address related but distinct analytical objectives. MDpocket extends the Fpocket algorithm to MD trajectories by computing pocket occupancy frequencies across frames on a fixed grid, thereby producing volumetric maps that are particularly useful for visualizing recurrent cavities and transient pocket formation. However, this grid-based representation differs from the centroid-based pocket descriptions generated by the tools included in our pipeline, making direct integration into a unified consensus framework non-trivial. Moreover, MDpocket derives persistence from the recurrence of spatial grid occupancy, whereas our framework estimates persistence from the agreement of multiple algorithmically diverse predictors applied independently to individual snapshots. These two strategies provide complementary views of pocket dynamics rather than directly interchangeable outputs. Similarly, TRAPP and related tools such as POVME and EPOCH are designed primarily for detailed characterization of predefined binding regions, including changes in pocket volume, shape, accessibility, and physicochemical properties along a trajectory. This makes them highly valuable when the binding site of interest is already known or can be reliably defined in advance. By contrast, the present framework was designed for de novo identification of candidate binding regions in conformational ensembles, without imposing prior spatial constraints. Accordingly, MDpocket, TRAPP, POVME, and EPOCH should not be viewed as competing approaches, but rather as complementary methods that can be applied downstream of a discovery-oriented workflow. In practice, the consensus pipeline described here may serve as an upstream step to identify persistent candidate pockets, which can then be subjected to more detailed dynamic characterization using tools such as MDpocket, TRAPP, or POVME.

#### P2Rank

P2Rank leverages a machine learning framework based on a Random Forest classifier to integrate a range of structural and physicochemical descriptors for the prediction of ligandable regions on the protein surface [26,41]. The algorithm begins by generating a uniform distribution of points across the solvent-accessible surface (SAS) of the protein, computed using the Connolly rolling-ball method. Each point is annotated with physicochemical features derived from the local environment of solvent-exposed atoms. These annotated points are then scored by the classifier to estimate their likelihood of being part of a ligand-binding site. High-scoring points are clustered spatially to form putative binding pockets, which are subsequently ranked according to their cumulative ligandability scores [38]. In this study, P2Rank v2.5.1 was executed with default parameters; all predicted pockets were retained without applying score-based filtering, to ensure unbiased sampling of candidate binding regions across the conformational ensemble. A characteristic feature of P2Rank lies in its ability to identify binding sites without reliance on predefined clustering thresholds, thereby allowing the detection of diverse pocket geometries [30]. Output files generated by P2Rank include a ranked list of predicted pockets with ligandability scores and centroid coordinates (predictions.csv), the protein residues associated with each pocket (residues.csv), PyMOL and ChimeraX visualization files, and the annotated SAS point set. For the downstream consensus analysis, pocket centroid coordinates were extracted from predictions.csv.

#### Fpocket

Fpocket is a geometry-based method that identifies potential ligand binding sites through the analysis of α-spheres, geometric entities derived via Voronoi tessellation. Introduced by Liang and Edelsbrunner [42], an α-sphere is defined as a sphere whose surface contacts exactly four protein atoms and contains no other atom in its interior [18]. Fpocket analyzes the spatial distribution of these α-spheres across the protein surface and aggregates them into clusters that are hypothesized to correspond to ligand-binding pockets. The clustering procedure is guided by three principal criteria: spatial proximity, topological connectivity, and minimum cluster size. Fpocket complements the geometric detection of cavities with structural and physicochemical descriptors, used in the subsequent characterization and prioritization of pockets.

Following the initial clustering, the algorithm applies geometric filters that exclude cavities too small to accommodate a ligand or completely buried and thus inaccessible to solvent [41]. The remaining pockets are then ranked using a composite druggability score that integrates geometric and physicochemical descriptors (e.g., polarity, hydrophobicity, mean α-sphere radius), prioritizing cavities with features statistically associated with known ligand-binding sites. Output files generated by Fpocket include a per-pocket descriptor file (_info.txt), a copy of the input PDB structure with α-sphere centers added as dummy atoms (_out.pdb), the α-sphere centers and radii of each pocket (_pockets.pqr, _vert.pqr), the protein atoms contacted by α-spheres in each pocket (_atm.pdb), and PyMOL and VMD visualization scripts.

In this study, Fpocket v3.0 was executed with default parameters; all predicted pockets were retained without applying score-based filtering, to ensure unbiased sampling of candidate binding regions across the conformational ensemble. For the downstream consensus analysis, pocket centroid coordinates were extracted from _info.txt.

#### Autosite

AutoSite is an energy-based method that identifies protein binding sites through the analysis of interaction energy distributions computed on the protein surface. By mapping the spatial distribution of energetically favorable protein–ligand interactions, AutoSite identifies surface regions compatible with ligand binding [22]. The core of the AutoSite algorithm involves the analysis of affinity maps generated by AutoDock, which model the interaction energies between the protein and three virtual probes: an aliphatic carbon probe (C, modeling hydrophobic interactions), a hydrogen-bond acceptor probe (OA), and a hydrogen-bond donor probe (HD). For each probe map, grid points with interaction energies below a probe-specific energy cutoff are retained as energetically favorable. The three filtered maps are then combined to retain only grid points where hydrophobic, hydrogen-bond acceptor, and hydrogen-bond donor interactions are simultaneously favorable. The resulting points are clustered spatially to define candidate binding pockets, and the most energetically stable point within each cluster is designated as a feature point. Candidate pockets are then filtered by minimum cluster size and ranked by an overall score that combines energy density and spatial compactness, prioritizing clusters with a high concentration of energetically favorable points within a compact volume.

AutoSite is natively integrated with the AutoDock framework, sharing its force field and grid representation for input map generation. In this study, AutoSite v1.0 was executed with default parameters using AutoGrid maps generated at 1.0 Å spacing for the C, OA, and HD probes; all predicted clusters were retained without score-based filtering, to ensure unbiased sampling of candidate binding regions across the conformational ensemble. Output files generated by AutoSite include one PDB file per cluster containing the cluster points (*_CL_*.pdb), one PDB file per cluster containing the feature points (*_fp_*.pdb), and a summary file reporting the scoring function values for each cluster (*_summary.csv). The AutoGrid input and output files (rigidReceptor.gpf, rigidReceptor.glg, rigidReceptor.x.map) provide the interaction energy maps used by AutoSite. For the downstream consensus analysis, the spatial coordinates of the cluster centers were extracted from *_CL_*.pdb.

#### EasyMIFs/SiteHound

EasyMIFs and SiteHound form a sequential energy-based pipeline for the identification of ligand-binding sites, in which EasyMIFs computes the protein interaction field and SiteHound detects binding sites from the resulting energy map.

EasyMIFs adopts an energy-based methodology, relying on Molecular Interaction Field (MIFs) analysis to detect regions of the protein that are energetically favorable for ligand interaction. The algorithm evaluates the interaction energies between the protein and a chosen molecular probe. In this study, the carbon probe CMET was exclusively employed, as it maps hydrophobic and van der Waals interactions, which dominate the contacts of most drug-like ligands and provide a broad definition of the ligand-occupiable volume. To refine the detection and classification of ligandable regions, an energy cutoff is applied to filter out grid points associated with unfavorable interactions [23]. SiteHound takes the EasyMIFs interaction energy map as input and identifies binding sites by clustering grid points with energetically favorable values. The clusters are ranked by their Total Interaction Energy (TIE), which sums the interaction energies of all grid points belonging to the cluster, prioritizing regions where energetically favorable interactions are spatially concentrated [23,25].

In this study, EasyMIFs and SiteHound v0.1 were executed with default parameters. EasyMIFs generates the interaction energy map (*.dx, in kJ/mol), which SiteHound processes to produce the set of predicted clusters: the points passing the energy filter (*.tmp), a per-cluster summary including Total Interaction Energy values (summary.dat), the points belonging to each cluster (clusters.dat), the residues in contact with each cluster (predicted.dat), and cluster representations for visualization (clusters.dx, clusters.pdb). The raw output of the SiteHound-EasyMIFs pipeline was subjected to a per-frame filtering protocol to remove low-intensity sites prior to inter-tool comparison. This step was necessary because the unfiltered predictions were considerably more numerous than those yielded by the other tools, making a direct comparison of the raw datasets methodologically unsound. To establish a principled filtering threshold, an optimal ‘elbow’ point within the interaction energy distribution was identified using the Kneedle algorithm [42], thereby retaining only sites above the inflection point of the energy curve and discarding marginal predictions. For the downstream consensus analysis, pocket centroid coordinates were extracted from the retained clusters (clusters.pdb).

#### DeepPocket

DeepPocket is a deep learning-based method that builds on Fpocket as an upstream candidate generator and refines its prediction through two sequential convolutional neural networks (CNNs) modules for both scoring and segmentation of binding sites [32]. The method begins by preprocessing protein structures followed by initial cavity detection using Fpocket. Each candidate center is voxelized into a 3D grid using libmolgrid [43], with atomic types represented as distinct input channels. In our protocol, we set the radius parameter r to 5 Å, defining a cubic grid with a side length of 10 Å at a resolution of 0.5 Å. This setting was adopted instead of the default (r=3 Å, box=6 Å) to enlarge the spatial context around each candidate pocket. The expanded box was selected to fully encapsulate larger or elongated cavities and to provide the CNN with a wider receptive field around each candidate center. In this study, DeepPocket was executed with the pre-trained classification and segmentation checkpoints provided by the authors. The number of ranked pockets was set to 15 (default: 1), the segmentation threshold to 0.01 (default: 0.5), and the residue-to-mask distance to 5.0 Å (default: 3.5 Å).

Output files generated by DeepPocket include the CNN-ranked list of predicted pockets, each represented by its barycenter coordinates (bary_centers_ranked.types), the corresponding CNN confidence scores (*_confidence.txt), and the volumetric segmentation masks produced by the U-Net module (one per predicted pocket). The pipeline also retains the Fpocket-derived intermediate files (*_out.pdb, *_pockets.pqr, visualization scripts), which correspond to the candidate pockets generated by Fpocket before CNN rescoring and segmentation. For the downstream consensus analysis, pocket barycenter coordinates were extracted from bary_centers_ranked.types.

#### SiteFerret

SiteFerret is a geometry-based method that identifies candidate binding sites through the clustering of spherical probes, sampled on the solvent-excluded surface (SES) of the protein at multiple probe radii (1.4-3.0 Å). The SES of the target protein is constructed using NanoShaper[44], and it is represented as a discrete point set, and sampling at multiple probe radii captures cavities of different sizes, from shallow grooves to deeper pockets. In SiteFerret, a cluster is designated as a conventional pocket if it comprises at least 100 probe spheres and terminates in a sequence of four or more aligned probes of increasing radius-a feature referred to as a pseudo-mouth. Subpockets are defined as minimal subclusters that satisfy these same size and alignment criteria, representing finger-grained concave subregions within larger cavities. Once the hierarchical clustering process yields a set of pockets and subpockets, SiteFerret characterizes each candidate through the integration of geometric and chemical properties. Candidate pockets are then scored using an unsupervised approach: it employs Isolation Forest, a classical anomaly detection algorithm, to score candidate pockets based on their deviation from the feature distribution of typical surface cavities.

In this study, SiteFerret was executed with default parameters. The PQR files generated by SiteFerret were processed using the loop.py script, which extracts the top 10 ranked pockets per structure. Because SiteFerret can produce a large number of candidate pockets, the analysis was restricted to the top 10 best-performing predictions per frame, consistent with the default output of the loop.py script and analogous to the per-frame filtering applied to the SiteHound-EasyMIFs pipeline. Output files generated by SiteFerret include a ranked summary of the predicted pockets (output_<PQR_NAME>.txt), and a dedicated folder (<STRUCTURE_NAME>_Pfiles/) containing per-pocket representations: the probe spheres of each cluster (clusterPocket<N>.pqr), the protein surface atoms within the pocket envelope (p<N>_atm.pqr), the pocket surface triangulation (p<N>.off), and a text file reporting the residues and pseudo-mouth information (infoPocket<N>.txt). Equivalent files are generated for the subpockets (with sub prefix), together with the NanoShaper surface triangulation (<STRUCTURE_NAME>.vert, <STRUCTURE_NAME> .face) and standard log files. For the downstream consensus analysis, pocket centroid coordinates were computed as the geometric center of the probe spheres stored in clusterPocket<N>.pqr.

### Benchmark dataset

The human glucose transporter 1 (GLUT1), a member of the major facilitator superfamily responsible for the facilitated diffusion of glucose across cell membranes, was selected as a representative model of a dynamic membrane transporter, owing to its well-characterized alternating-access mechanism and the availability of multiple structural states of its transport cycle.

Outward-open and outward-occluded conformations were generated by homology modeling using GLUT3 templates (PDB IDs: 4ZWB and 4ZW9), while the crystal-resolved inward-open configuration (PDB ID: 4PYP) was employed as the final target structure for MD simulation. The system was embedded in a POPC lipid bilayer using CHARMM-GUI, solvated with TIP3P water, and neutralized with Na□/Cl□ ions. A hybrid Supervised Molecular Dynamics (SuMD) [46] and Targeted Molecular Dynamics (TMD) protocol was employed to investigate glucose transport dynamics and GLUT1 conformational transition. In brief, SuMD was used to guide D-glucose into the central binding site, while TMD was applied to simulate conformational transitions between outward-open, outward-occluded, and inward-open states.

After energy-minimization and equilibration, ten independent MD simulations were performed using the Amber ff14SB force field in Amber 22 [47]. SuMD simulations were conducted in short adaptive intervals (300 ps), followed by TMD runs (10 ns each) to induce conformational transitions and ligand translocation toward the cytoplasmic side. NetMD [48] was used to evaluate the consensus between replicas and the most representative one was used for the binding site prediction analyses, using a total of 500 frames. To better capture the conformational variability of the protein, the trajectory was partitioned into two distinct phases according to the structural transitions observed. The first phase comprised frames 1-100, in which the protein was guided from the *outward-open* to the *outward-occluded* conformations, while the second phase included frames 101-500, where the structure shifted from *outward-occluded* to *inward-open* configuration. Dividing the trajectory into these phases allowed for a more refined analysis, enabling the monitoring of cryptic pockets emerging during conformational changes as well as the identification of binding sites that remain conserved throughout the simulation.

To further validate the workflow on a well-characterized cryptic system, the human Aldose Reductase (AR) was included as reference. AR is a paradigmatic model of structural plasticity, known for the opening of a “*specificity pocket*” mediated by the Leu300 loop movement [35,36]. The inclusion of the AR set allowed for testing algorithm sensitivity and the ability to identify transient pockets. For the AR analysis, we utilized the molecular dynamics data provided by the TRAnsient Pockets in Proteins (TRAPP) portal, which includes 59 conformational frames. On this dataset, we performed the same prediction and consensus analyses as those conducted for GLUT1, ensuring methodological consistency between the two studied systems.

### Statistical correlation framework

To quantify the inter-tool concordance of pocket predictions across the conformational ensemble, we performed a Pearson correlation analysis. To enable a residue-level comparison across tools, we computed for each frame and residue the Euclidean distance between the residue geometric center and the centroid of the nearest pocket predicted by each tool Pearson correlation coefficients were then computed between the per-residue distance vectors of each tool pair, providing a quantitative measure of inter-tool spatial agreement across the full residue set. Pearson correlation was selected over rank-based alternatives given the continuous and approximately linear nature of the distance distributions observed across tools.

### Methodological framework for consensus and concordance analysis

Several metrics have been proposed to compare predicted binding pockets, including distance-based and overlap-based measures. Among the most commonly used are the distance to closest centroid (DCC), which measures the minimal distance between predicted and reference pocket centers [49,50]; the Discrete Volume Overlap (DVO), which compares shapes of the predicted and actual pockets; the distance to closest atom (DCA), the distance from the predicted pocket center to the nearest ligand atom [49,50]; and the Overlap (OLP) score or Overlap ratio (RO), which assesses the extent of common residues between predicted and experimental binding sites [49][51]. However, their direct applicability is limited by the heterogeneous nature of outputs produced by the selected tools, which range from geometric descriptors (Fpocket, SiteFerret) to energy-based affinity grids (AutoSite, SiteHound/EasyMIFs) and volumetric segmentation masks (DeepPocket). To address this limitation and ensure a robust comparison across heterogeneous tool outputs, we implemented a dual consensus framework combining HDBSCAN-based spatial clustering of pocket centroids and volumetric comparison based on the Intersection over Union (IoU) metric. The two components of the framework were designed to operate independently and on distinct geometric principles, so that their agreement constitutes a form of cross-validation rather than redundancy. A binding site supported by both methods (HDBSCAN clustering and IoU analysis) is therefore identified not solely as a spatially recurrent region, but as one that is simultaneously coherent in its centroid-level spatial distribution and consistent in its three-dimensional volumetric architecture across heterogeneous algorithmic outputs. This dual requirement effectively filters out noise while preserving genuine consensus sites across methodologically diverse tools.

#### Clustering methods

To identify spatially coherent binding regions across the MD ensemble, we employed a clustering strategy based on pocket centroids pooled across all frames and tools. Two density-based clustering algorithms were tested: DBSCAN (Density-Based Spatial Clustering of Applications with Noise) [44,52] and HDBSCAN (Hierarchical DBSCAN) [53]. DBSCAN is a density-based method that groups closely packed points (requiring only two parameters: ε (neighborhood radius and minPts).

### HDBSCAN

HDBSCAN is an extension of DBSCAN that builds a *hierarchy* of clusters at different density levels and then selects the most stable clusters adapting its density threshold locally rather than applying a single global ε. This makes it particularly suited to pocket detection across MD ensembles, where some binding sites recur frequently and form dense point clouds while others appear only transiently. The key parameter controlling cluster granularity is min_cluster_size, which was initialized at 50% of the total number of frames per phase as a baseline threshold for pocket persistence, and subsequently refined empirically for each tool based on the spatial coherence and stability of the resulting clusters (*See Table 1 Supp. Tables*). In both cases, points failing to meet the minimum density criteria are labeled as outliers.

#### IoU

To provide an independent volumetric validation of the HDBSCAN-based spatial clustering, we implemented an alternative consensus strategy based on the Intersection over Union (IoU) metric, also known as the Jaccard Index. This approach serves as an independent validation metric, shifting the focus from centroid proximity to a more detailed assessment of volumetric overlap. To standardize the comparison across heterogeneous tool outputs, all predicted binding sites were projected onto a common volumetric representation. For each method, a pocket was defined as a three-dimensional binding region obtained by aggregating its predicted points. The binding space was discretized onto a voxel grid with a resolution of 1 Å³, and a voxel was considered occupied if it lay within a cutoff radius (R= 5 Å) [54,55] of any pocket-defining element (e.g. a-spheres for geometry-based tools, energy grid points for energy-based tools and atomic coordinates for machine/deep learning-based tools) generated by the method. For every MD snapshot, the spatial agreement between two pockets prediction A and B was quantified using the IoU metric, defined as:

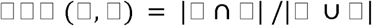

where |□ ∩ □| represents the number of shared occupied voxels and |□ ∪ □| the total unique occupied voxels associated with the two methods. An IoU threshold greater than 0.5 was adopted to define a significant consensus between pocket predictions [56].

#### Persistence frequency

We computed temporal persistence to evaluate both the stability and the recurrence of consensus pockets across the MD trajectory. For each consensus cluster, persistence was computed independently for each tool contributing to that cluster and then averaged across tools, with the standard deviation reported as a measure of inter-tool variability. For a given tool and cluster, the detected frames were grouped into contiguous runs, where two consecutive detections were considered part of the same run if the gap between them did not exceed a defined tolerance δ. The length of each run was defined as an interval from the first to the last detected frame, inclusive of internal gaps. Only runs meeting a minimum length threshold of 20% of the total frames in each phase (20 frames in phase 1; 80 frames in phase 2) were considered valid, in order to filter out spurious short-lived detections caused by local side-chain rearrangements or transient solvent-accessible cavities [45][57]. Per-tool persistence was then computed as the sum of valid run lengths divided by the total number of frames in the phase, expressed as a percentage.

To capture the full spectrum of pocket dynamics, ranging from constitutively accessible sites to transiently sampled regions, two persistence criteria were defined based on the gap tolerance parameter δ. Under the Strict Persistence criterion (δ = 0), no gaps were tolerated between detections: only perfectly consecutive frames contributed to a valid run. Under the Soft Persistence criterion (δ = 5), runs separated by up to five missing frames were merged, accommodating brief detection discontinuities arising from local structural fluctuations or tool-specific sensitivity limits. The gap tolerance parameter δ was set to 5 frames based on a sensitivity analysis performed across increasing values of δ (*See Fig. S2 Supp. Materials*). For the HDBSCAN-based persistence, values remained substantially stable across the tested range, with no meaningful change in per-cluster persistence scores beyond δ = 5. For the IoU-based consensus, persistence values were unaffected by δ by construction, as the underlying volumetric clustering does not depend on frame-level detection continuity. The value of δ = 5 was therefore selected as the minimum sufficient to absorb brief detection discontinuities without inflating persistence estimates. Based on MDpocket criteria, an occurrence frequency of 20% is associated with druggable binding sites[45]; accordingly, a Strict Persistence threshold of 20% was adopted to identify stable, recurrently accessible pockets. A Soft Persistence threshold of 10% was additionally applied to capture transiently sampled sites that are recurrently visited but not continuously maintained across the trajectory.

## RESULTS

### Inter-tool consistency

#### GLUT1

Concordance analysis revealed a highly variable landscape with modest peaks, where the degree of agreement between different tools varied systematically as a function of the methodological principles underlying each tool. The highest correlation value was observed between Fpocket and AutoSite (Pearson correlation coefficient, r = 0.363) when analyzing the GLUT1 conformational ensemble (**Table 2**); this is a particularly informative result given that these two tools rely on fundamentally different principles, i.e., geometric cavity detection and energy-based affinity mapping, respectively, suggesting that, despite their methodological divergence, both approaches converge in identifying regions that are simultaneously geometrically accessible and energetically favorable for ligand binding. Comparable levels of agreement were found between P2Rank and AutoSite (r = 0.355) and between SiteHound and AutoSite (r = 0.327), indicating that AutoSite predictions tend to be broadly consistent across the different algorithmic families included in this study. Conversely, the lowest correlation was recorded between SiteFerret and DeepPocket (r = 0.053), highlighting how deep learning-based volumetric representations can diverge substantially from geometry-based descriptors, likely reflecting fundamentally different feature spaces used to characterize protein surface topology. Notably, DeepPocket showed consistently low agreement with most other tools, with Pearson correlation coefficients ranging from 0.053 to 0.274, suggesting that its learned representations capture pocket features that are complementary to those captured by geometry- or energy-driven descriptors.

**Table 2.** Pearson correlation matrix to quantify inter-tool agreement for the GLUT1 conformational set. The matrix displays the correlation coefficients between the minimum residue-to-pocket distances calculated by six detection algorithms.

|  | AutoSite | DeepPocket | Fpocket | P2rank | SiteFerret | SiteHound |
| --- | --- | --- | --- | --- | --- | --- |
| AutoSite | 1.000 | 0.195 | 0.363 | 0.355 | 0.303 | 0.327 |
| DeepPocket | 0.195 | 1.000 | 0.181 | 0.274 | 0.053 | 0.094 |
| Fpocket | 0.363 | 0.181 | 1.000 | 0.180 | 0.242 | 0.093 |
| P2rank | 0.355 | 0.274 | 0.180 | 1.000 | 0.173 | 0.320 |
| SiteFerret | 0.303 | 0.053 | 0.242 | 0.173 | 1.000 | 0.159 |
| SiteHound | 0.327 | 0.094 | 0.093 | 0.320 | 0.159 | 1.000 |

Taken together, these results indicate that AutoSite occupies a central position in the inter-tool correlation landscape, acting as a methodological bridge across algorithmic families, while DeepPocket consistently occupies the periphery, capturing a distinct and complementary view of the binding surface

#### AR

Compared to the GLUT1 ensemble, the AR conformational set (**Table 3**) showed a marked and consistent increase in inter-tool concordance, with higher correlation values observed across all tool pairs.

**Table 3.** Agreement between tools for the AR conformational set assessed by Pearson correlation. The matrix displays the correlation coefficients (*r*) between the minimum residue-to-pocket distances calculated by six detection algorithms.

|  | AutoSite | DeepPocket | Fpocket | P2rank | SiteFerret | SiteHound |
| --- | --- | --- | --- | --- | --- | --- |
| AutoSite | 1.000 | 0.516 | 0.425 | 0.394 | 0.414 | 0.474 |
| DeepPocket | 0.516 | 1.000 | 0.415 | 0.573 | 0.444 | 0.590 |
| Fpocket | 0.425 | 0.415 | 1.000 | 0.304 | 0.394 | 0.336 |
| P2rank | 0.394 | 0.573 | 0.304 | 1.000 | 0.352 | 0.489 |
| SiteFerret | 0.414 | 0.444 | 0.394 | 0.352 | 1.000 | 0.385 |
| SiteHound | 0.474 | 0.590 | 0.336 | 0.489 | 0.385 | 1.000 |

Within the AR ensemble, correlations frequently exceed 0.5, with particularly striking peaks between DeepPocket and SiteHound (r = 0.590) and DeepPocket and P2rank (r = 0.573), values that stand in sharp contrast with the consistently low DeepPocket correlations observed with GLUT1.

Notably, this convergence is most pronounced for DeepPocket, which showed the lowest pairwise correlation coefficients across all tool pairs in GLUT1 (0.053-0.274) but that reaches the two highest values in AR (r = 0.590 with SiteHound, r = 0.573 with P2Rank). This result suggests that the degree of agreement between different methodological classes may be influenced by the underlying conformational landscape.

### Binding Site Consensus Analysis: Pocket Prediction, Dataset Construction

Application of the six selected tools to the GLUT1 MD trajectory (see Methods) generated an extensive dataset of predicted pocket locations spanning both its conformational phases, outward-open (phase 1) and outward-occluded (phase 2).

In phase 1, a total of 7,836 pockets were predicted, with individual contributions as follows: SiteFerret (1,000), AutoSite (1,556), Fpocket (2,989), P2Rank (1,078), SiteHound-EasyMIF (1,213; filtered via the Kneedle algorithm to remove low-intensity sites), P2Rank (1,078), DeepPocket (1,053). In phase 2, the number of pockets increased substantially, reaching a total of 35,355 pockets (Fpocket: 12,536; AutoSite: 5,640; SiteHound/EasyMIFs: 4,702; P2Rank: 4,386; DeepPocket: 4,271; SiteFerret: 4,000), representing an approximately 4.5-fold increase over phase 1, which is consistent with the greater conformational heterogeneity of the inward-open state.

For P2Rank and DeepPocket, all predicted cavities were retained regardless of their internal confidence scores. This choice ensures unbiased sampling of the conformational ensemble, capturing transient pockets that might receive lower scores in static context but demonstrate significant persistence or consensus within a dynamics ensemble (see Methods).

#### Clustering of detected pockets

To group recurring predictions into spatially coherent binding regions, density-based clustering was applied to the centroid locations pooled across all frames and tools. Clustering outcomes varied depending on the tool and the conformational phase of the protein. Across all six tools and both phases, HDBSCAN consistently produced a more compact set of consensus clusters compared to DBSCAN, which instead generated a larger number of smaller groups and a higher proportion of outlier points (cluster -1, **Figure 1**). This difference was particularly pronounced for tools producing dense and spatially extended outputs such as DeepPocket and P2Rank, where DBSCAN fragmented the predictions into a substantially larger number of clusters than HDBSCAN, suggesting a greater sensitivity to local density variations. Conversely, geometry-based tools (Fpocket, SiteFerret) showed comparable cluster counts under the two methods, indicating that their outputs are intrinsically more spatially compact and less affected by the clustering algorithm. A general trend toward increased cluster numbers in phase 2 was observed for most tools, consistent with the broader conformational sampling and the higher spatial dispersion of predicted sites associated with the inward-open state. This behavior is illustrated in **Figure 1**, which shows the comparison between HDBSCAN and DBSCAN clustering for AutoSite as a representative case. The variability observed between DBSCAN and HDBSCAN-derived cluster counts highlighted the need to assess the relative strengths of the two approaches in order to select the most appropriate method for further analysis.

**Figure 1.**
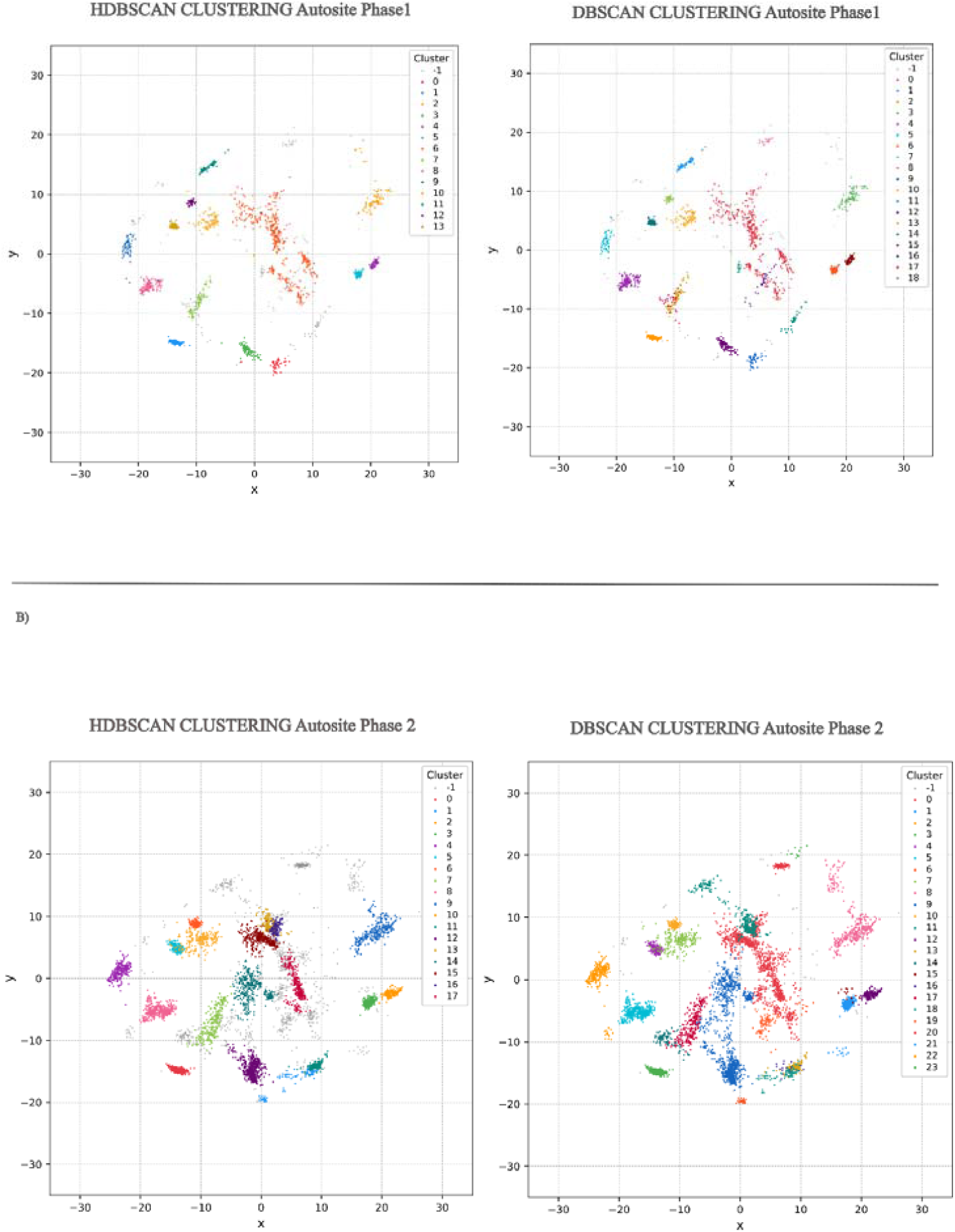
Comparison of HDBSCAN and DBSCAN clustering applied to AutoSite-identified pockets. A) phase 1: HDBSCAN yields 14 clusters, DBSCAN produces 19, with greater fragmentation and more outliers (cluster -1). B) phase 2: HDBSCAN yields 18 clusters, DBSCAN produces 24.

To validate clustering on a structurally defined ground truth, HDBSCAN was applied to the pocket centroids of the AR conformational ensemble. The number of clusters identified varied substantially across tools, reflecting differences in detection sensitivity and spatial resolution (*See Table 2 Supp. Tables*).

SiteHound produced the highest number of clusters (48), reflecting the high sensitivity of this energy-based approach to local surface fluctuations, which generates a large number of spatially distinct but short-lived sites. In contrast, P2Rank yielded the most compact clustering output (5 clusters), consistent with its machine learning-based approach that prioritizes high-confidence, well-defined binding regions. Fpocket (17), AutoSite (13), DeepPocket (8), and SiteFerret (7) yielded intermediate counts, thereby reflecting the different trade-offs between sensitivity and specificity inherent to each methodological category. This tool-specific variability in cluster granularity underscores the complementary nature of the six detection strategies and directly motivates the consensus-based integration approach described in the following section.

### Identification of Consensus Pockets

#### Consensus Intra-Category GLUT1

To assess the internal consistency of pocket predictions among tools operating on similar methodological principles, we performed an intra-category consensus analysis applying the 6.5 Å centroid-to-centroid threshold (*See Supp. Materials*) independently of each methodological family (geometry-based, energy-based, and machine/deep learning) and to each conformational phase (**Figure 3**).

**Figure 3.**
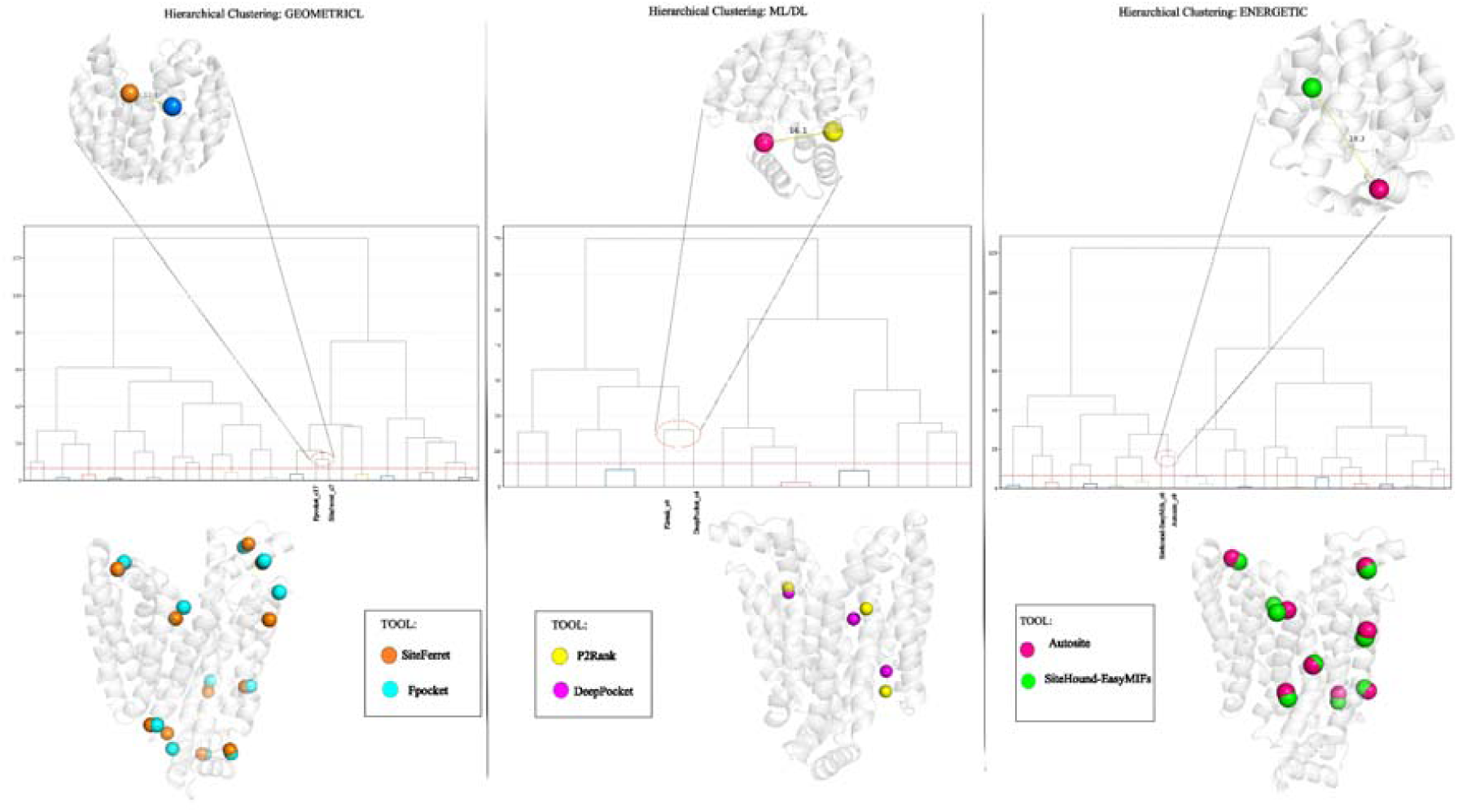
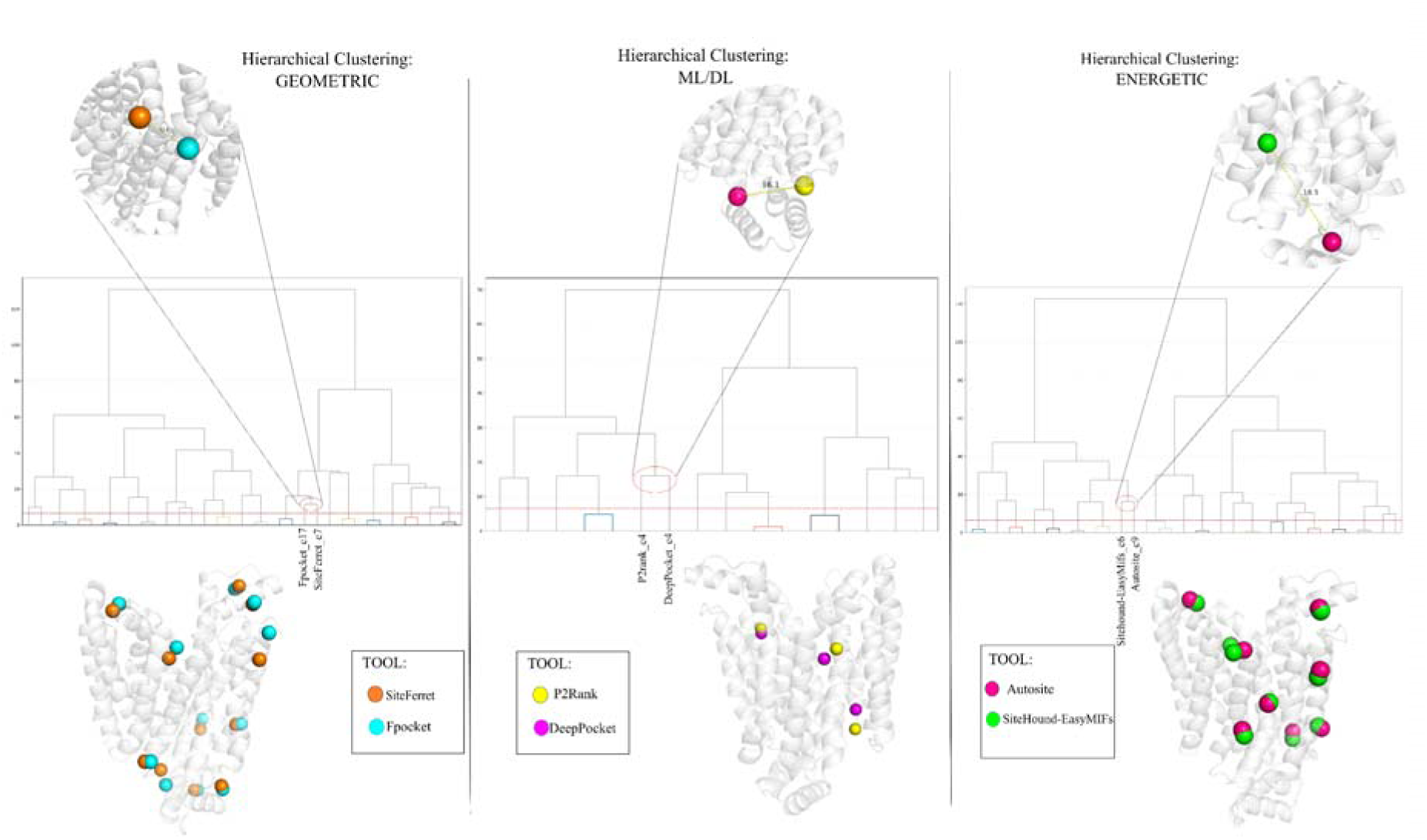
Intra-category hierarchical clustering results for phase 1, organized by methodological category: geometric (left, Fpocket and SiteFerret), ML/DL (center, P2Rank and DeepPocket), and energetic (right, AutoSite and SiteHound-EasyMIFs) The vertical axis represents the Euclidean distance (in Å) at which pairs of clusters or cluster groups are merged during the agglomerative clustering procedure, with greater heights indicating that the joined elements are spatially farther apart. The dashed red line marks the 6.5 Å distance threshold used to define consensus groups: only centroids merged below this height are considered part of the same consensus cluster. The inset panels highlight representative cases in which two clusters from different tools exceed this threshold and are therefore not merged into a single consensus: Fpocket_c17 and SiteFerret_c7 (11.45 Å, geometric); P2Rank_c4 and DeepPocket_c4 (16.06 Å, ML/DL); SiteHound_c6 and AutoSite_c9 (18.32 Å, energetic). The bottom row shows the three-dimensional distribution of all matched consensus centroids mapped onto the GLUT1 structure in phase 1, with spheres colored by tool of origin.

##### Geometric category (Fpocket - SiteFerret)

In phase 1, Fpocket and SiteFerret yielded a total of 10 consensus clusters within the 6.5 Å threshold. The matched clusters span multiple structurally distinct regions of the protein surface, indicating spatial agreement between the two geometry-based tools across the phase 1 conformational ensemble. In phase 2, the number of total geometric consensus clusters increases to 14, reflecting a greater spatial overlap between Fpocket and SiteFerret in this conformational state. This increase suggests that the phase 2 ensemble exposes a larger number of geometric cavities that are simultaneously detectable by both tools.

##### Energetic category (AutoSite - SiteHound)

In phase 1, AutoSite and SiteHound produced 9 matched pairs. The matched clusters cover a broad range of protein regions, consistent with the sensitivity of energy-based tools to electrostatic and hydrophobic surface properties. In phase 2, the total number of energetic consensus clusters rises to 13, representing the highest absolute number of intra-category matches across all three categories in phase 2. The increase in matched pairs from phase 1 to 2 suggests enhanced convergence between AutoSite and SiteHound in the phase 2 conformational state, potentially reflecting the emergence of energetically favorable cavities in the inward-open conformation.

##### ML/DL category (P2Rank - DeepPocket)

In phase 1, P2Rank and DeepPocket agreed only on 3 clusters. This was the lowest consensus cluster count among the three methodological families in phase 1, reflecting the fundamental structural differences between a surface-point Random Forest classifier (P2Rank) and a volumetric CNN segmentation approach (DeepPocket), which operate on distinct spatial representations of the protein. In phase 2, the total number of ML/DL consensus clusters increased to 4, the smallest absolute gain among the compared classes. The limited overlap between P2Rank and DeepPocket in both phases is consistent with their distinct training objectives and feature representations, which may lead to partial but not complete spatial convergence.

Across all three categories, a consistent pattern emerged: the number of matched pairs increases from phase 1 to phase 2, with the energetic category showing the greatest absolute gain. Although phase 2 contains substantially more simulation frames than phase 1 (400 vs. 100), the observed expansion is consistent across all three methodological categories and reflects the broader conformational diversity of the inward-open state sampled in phase 2 compared to the outward-open to outward-occluded transition of phase 1.

To assess whether these intra-category patterns extend to a system with a known cryptic pocket, the same analysis was applied to the AR conformational ensemble.

#### Consensus Intra-Category AR

The intra-category consensus analysis was also applied to the AR conformational ensemble. The known specificity pocket is defined by the C-terminal loop residues Ala299-Cys303, with Leu300 being the primary gatekeeper and Cys298 lining the adjacent hydrophobic cleft. The three methodological categories showed markedly different detection success. Energy-based tools (AutoSite, SiteHound) showed the highest detection accuracy, identifying a consensus pocket (*cluster 7*) sharing four residues with the specificity pocket (Cys298, Ala299, Leu300, Leu301). This reflects their ability to detect favorable probe-protein interaction potential even before full cavity formation, a key advantage for cryptic sites driven by transient loop displacement. ML/DL-based tools (P2Rank, DeepPocket) identified a consensus pocket (*cluster 3*) sharing only Cys298 with the target site, with remaining residues displaced relative to the known specificity pocket’s residues. This partial detection likely stems from training bias toward stable, well-defined binding sites, limiting sensitivity to the dynamically sampled conformations of this transiently accessible cryptic pocket. Geometry-based tools (Fpocket, SiteFerret) found no residue overlap with the specificity pocket, as they require pre-existing surface concavities and cannot sense interaction potential in the absence of a fully formed cavity (*See Fig. S3 Supp. Materials*).

#### Consensus Inter-Category (GLUT1)

To identify spatially recurring binding regions supported by tools spanning distinct methodological families, we performed an inter-category consensus analysis applying a selection criterion of at least four out of six tools (spanning geometry-based, energy-based, and machine/deep learning approaches) and a 6.5 Å inter-centroid distance threshold, with cluster numbering assigned independently for each phase.

#### Phase 1

In phase 1, this approach yielded 9 high-confidence consensus clusters. All nine clusters were supported by either 4 or 5 tools, with the majority (6 out of 9) reaching 5-tool agreement. Two clusters (*clusters 22* and *23*) were the only ones in which DeepPocket contributed, and *cluster 23* was the sole cluster spanning all three methodological categories (geometry-based, energy-based, and ML/DL), representing the highest possible inter-category convergence criterion. The remaining clusters were all supported by combinations of AutoSite, Fpocket, P2Rank, SiteFerret, and SiteHound with DeepPocket absent in all cases. The temporal stability of these regions is consistent with their spatial robustness: soft persistence values are uniformly high across all 9 clusters, ranging from 80.0% (*cluster 14*) to 100% (*cluster 29*), with inter-tool standard deviations remaining below 13.0% for six out of nine clusters.

Strict persistence values were more variable (range: 13.8% for *cluster 26* to 80.2% for *cluster 29*) reflecting differences in frame-by-frame detection stringency, but the gap between strict and soft remained moderate for most clusters, indicating that detections were largely continuous rather than episodic. *Cluster 29* stood out as the most stable site overall, achieving 100% soft persistence with zero inter-tool variability, while *cluster 14* showed the greatest heterogeneity with standard deviations of 49.0% and 40.0% in strict and soft mode respectively, suggesting substantial differences in frame-by-frame detection stringency across tools for this site. Overall, the phase 1 inter-category consensus landscape is characterized by high spatial robustness and uniform temporal stability, consistent with the relatively constrained conformational dynamics of the outward-open to outward-occluded transition.

#### Phase 2

In phase 2, the same criteria yielded 11 consensus clusters, reflecting a more fragmented pocket landscape. Five clusters reached 5-tool agreement and one, c*luster 36*, was supported by all six tools simultaneously, representing the highest inter-method convergence observed across both phases. Notably, SiteHound was absent from several phase 2 clusters where it contributed in phase 1, suggesting reduced detection sensitivity of this tool for the more transient cavities characterizing the phase 2 conformational ensemble. DeepPocket contributed only to *cluster 36* and *cluster 15*, confirming that ML/DL-based detection remained selective across both phases, consistently contributing to only the most robust consensus sites. The temporal profile of phase 2 clusters was substantially different from phase 1: soft persistence values spanned a much wider range, from 22.2% (*cluster 16*) to 94.4% (*cluster 36*), and inter-tool standard deviations were considerably higher across the board, frequently exceeding 25%. The pronounced dissociation between strict and soft modes is particularly informative: the complete absence of frame-by-frame strict detection for several clusters indicated that these sites were never continuously occupied across consecutive frames, while the appreciable soft persistence confirmed that they are nonetheless recurrently visited throughout the trajectory. This pattern is the hallmark of transient binding sites, pockets that open and close dynamically, accessible for discrete windows of time rather than maintained as stable cavities. The fact that such sites are captured by consensus across at least 4 independent tools further excludes stochastic noise as an explanation, supporting their classification as genuine but transiently sampled druggable regions.

Taken together, these results highlight a clear difference in the temporal character of the two phases: phase 1 was dominated by stable, persistently detected binding regions with high and consistent soft persistence, while phase 2 was characterized by a more heterogeneous ensemble of sites, several of which are only transiently accessible. This transition reflects the distinct conformational dynamics of GLUT1 across the two trajectory segments, corresponding to different stages of its alternating-access transport cycle.

#### Consensus Inter-Category (AR)

Extending the consensus analysis to the inter-category level, the same 4-out-of-6-tool protocol used to GLUT1 was applied to the AR conformational ensemble. This analysis identified two consensus regions in the vicinity of the known specificity pocket of AR, defined by the Leu300 loop rearrangement. Notably, *cluster 7* shared one residue with the target site (Cys298) reinforcing its relevance as a partial but consistent predictor of the cryptic site. *Cluster 4*, while not sharing any residue with the specificity pocket, is spatially proximal to the Leu300 loop region, suggesting that the inter-category consensus localized to the correct area of the protein surface even in the absence of exact residue overlap. This spatial convergence provided a meaningful validation of the computational workflow and, crucially, demonstrated how inter-category consensus can serve as an effective strategy to balance the complementary strengths and limitations of individual tools.

To evaluate the temporal stability of these pockets, we applied the dual persistence framework. Given the inherently transient nature of the Leu300-driven specificity pocket, our evaluation focused on Soft Persistence, which allows gaps of up to 5 frames and thus provides a more accurate quantification of biological recurrence across the 59-frame trajectory than Strict Persistence. Both pockets exhibited a robust temporal profile, justifying their classification as high-confidence sites:

- *Cluster 7* emerged as the most significant region, with a Soft Mean persistence of 69.5%, indicating that the site remained a recurring feature of the conformational landscape, despite the discontinuous nature of its detection due to loop flexibility.
- *Cluster 4* demonstrated comparable stability, with a Soft Mean of 71.5%.

Together, these results demonstrate that the combination of inter-category spatial consensus and soft temporal persistence successfully recovers and characterizes a well-documented cryptic pocket, providing a direct validation of the framework on a system with an experimentally established transient binding site.

### Alternative Consensus Strategy: IoU Analysis across Phases

To complement the spatial clustering of pocket coordinates, we applied the IoU method, as an independent volumetric consensus strategy. This approach provided an independent layer of evidence, testing whether the identified binding sites share reproducible 3D architectures across heterogeneous algorithmic principles. To obtain high-confidence predictions, the analysis was restricted to validated pockets that met a stringent dual filter: detection by at least four tools and a minimum IoU occupancy (fraction of frames above the IoU threshold) of 20%.

#### GLUT1 phase 1

To further validate the spatial consistency of the identified consensus regions, we compared the centroid positions of the HDBSCAN consensus clusters with those of the IoU trajectory clusters (Figure 4), computing the Euclidean distance between each matched pair. The matches were established based on spatial proximity and shared contributing tools, and the resulting centroid distances provided a quantitative measure of inter-method concordance.

**Figure 4.**
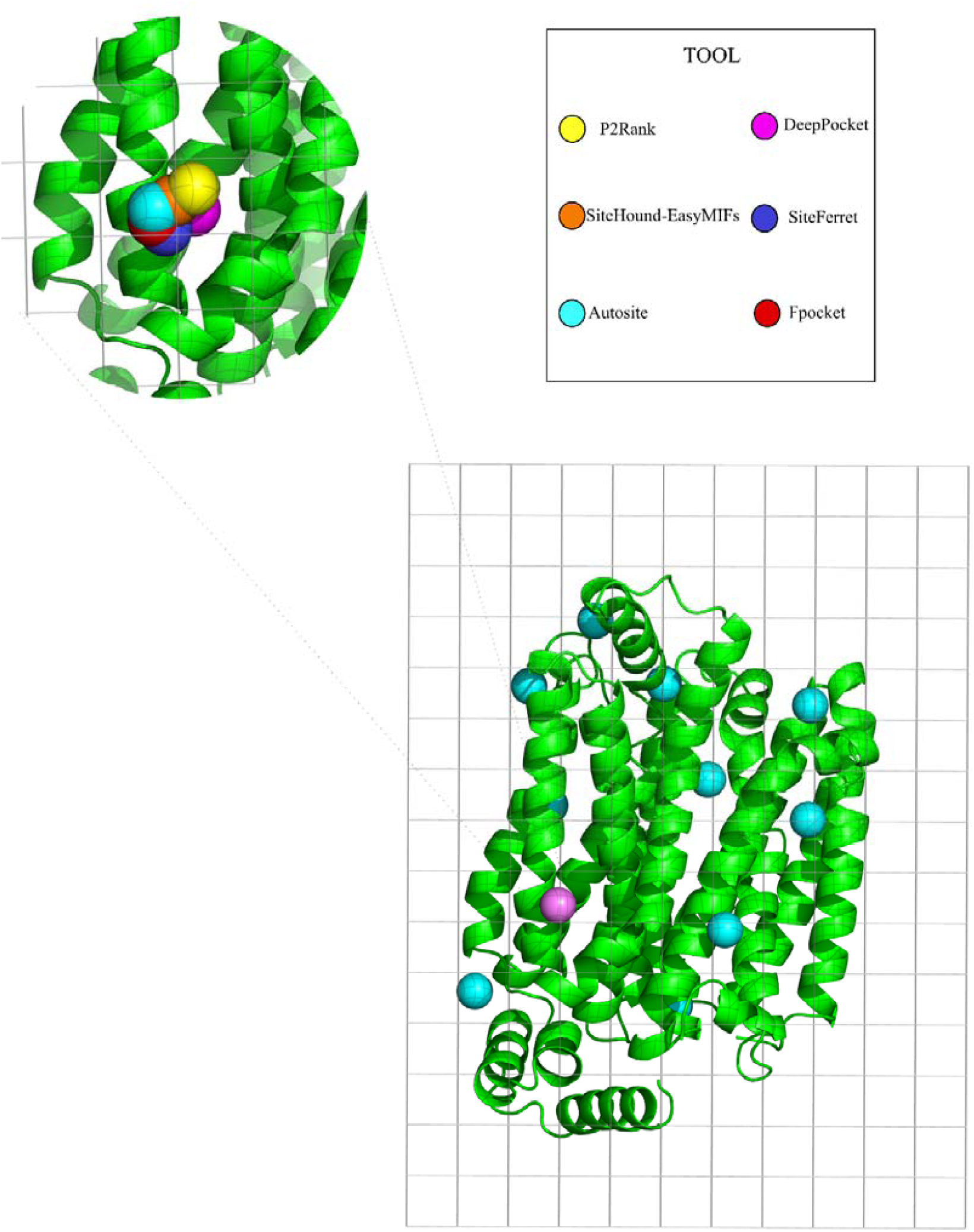
Schematic illustration of the IoU (Intersection over Union) grid-based method applied to pocket trajectory analysis. The main panel (bottom right) shows the full GLUT1 structure in phase 2 with all trajectory clusters identified by the IoU method mapped onto the protein surface. The inset (top left) provides a view of IoU trajectory 2, illustrating the spatial superposition of tool-specific pocket centroids within a defined grid cell: the overlapping spheres from six distinct tools demonstrate the high degree of spatial convergence that underlies the IoU consensus assignment for this cluster.

In phase 1, six HDBSCAN-IoU pairs were identified:

– *Cluster 14* vs *IoU 1* (distance: 0.33 Å): both methods detect a persistent site (IoU occupancy 100%, 5 tools: AutoSite, Fpocket, P2Rank, SiteFerret, SiteHound) in the same spatial region, with near-identical centroids.
– *Cluster 15* vs *IoU 4* (distance: 0.72 Å): IoU occupancy 77%, 5 tools. Spatially well-defined and consistently detected.
– *Cluster 23* vs *IoU 3* (distance: 0.76 Å): IoU occupancy 100%, 5 tools (AutoSite, DeepPocket, Fpocket, P2Rank, SiteHound). The only match in phase 1 where DeepPocket contributes in both methods, and the only HDBSCAN cluster spanning all three methodological categories.
– *Cluster 29* vs *IoU 11* (distance: 0.57 Å): IoU occupancy 26%, 5 tools. Despite lower IoU occupancy, this site corresponds to the most persistent HDBSCAN cluster in phase 1(soft 100% ± 0.0), suggesting that the IoU method captures it as a recurrent but shorter-lived trajectory.
– *Cluster 22* vs *IoU 19* (distance: 1.51 Å): IoU occupancy 21%, 5 tools (AutoSite, DeepPocket, Fpocket, SiteFerret, SiteHound). The second match where DeepPocket contributes in both methods.
– *Cluster 26* vs *IoU 9* (distance: 1.64 Å): IoU occupancy 28%, 5 tools. The largest centroid distance in phase 1, yet still well within the clustering threshold.

All six phase 1 pairs show centroid distances below 2 Å, confirming excellent spatial concordance between the two independent methods across the entire set of matched sites.

#### GLUT1 phase 2

In phase 2, six HDBSCAN–IoU pairs were identified:

- *Cluster 15* vs IoU 1 (distance: 0.23 Å): IoU occupancy 100%, 5 tools (AutoSite, DeepPocket, Fpocket, P2Rank, SiteHound). The closest centroid match in phase 2.
- *Cluster 36* vs IoU 2 (distance: 0.73 Å): IoU occupancy 100%, 6 tools, the only match in either phase where both methods achieve full 6-tool agreement simultaneously.
- *Cluster 7* vs IoU 3 (distance: 0.66 Å): IoU occupancy 60.75%, 5 tools (AutoSite, Fpocket, P2Rank, SiteFerret, SiteHound).
- *Cluster 26* vs IoU 4 (distance: 0.40 Å): IoU occupancy 26.25%, 5 tools.*Cluster 12* vs IoU 5 (distance: 1.93 Å): IoU occupancy 25.5%, 5 tools.
- *Cluster 5* vs IoU 8 (distance: 1.27 Å): IoU occupancy 22.75%, 5 tools.

As in phase 1, all phase 2 centroid distances fall below 2 Å, confirming robust spatial agreement between HDBSCAN and IoU. A notable pattern emerges when comparing the two phases: in phase 1, high IoU occupancy (77–100%) consistently corresponds to high HDBSCAN soft persistence, while low IoU occupancy sites (21–28%) are nonetheless captured with high soft persistence by HDBSCAN (91–97%), suggesting that the HDBSCAN framework integrates transient signals more effectively than the IoU trajectory-based approach. In phase 2, this divergence is more pronounced: two sites reach 100% IoU occupancy while HDBSCAN soft persistence ranges from 80 to 94%, and the remaining four sites show low IoU occupancy (22–26%) paired with highly variable HDBSCAN persistence (66–87%), reflecting the more heterogeneous and dynamic character of the phase 2 conformational ensemble.

## DISCUSSION

The identification of druggable binding sites remains a central challenge in structure-based drug discovery, particularly relevant for virtual screening protocols and for integrative docking approaches such as HADDOCK [58], where the definition of binding regions can be guided by experimental or predicted restraints. The proliferation of pocket prediction tools has not resolved, but rather amplified, the difficulty of making principled methodological choices. Most available tools were designed and benchmarked on static crystal structures, where pocket geometry is fixed and detection performance assessed against validated ligand positions. When applied to conformational ensembles derived from molecular dynamics simulations, however, this static-structure paradigm breaks down, with pockets switching between open and closed states, and the same protein region may be simultaneously detected by one algorithm and missed by another. In this dynamic context, inter-tool agreement represents a more informative metric than individual performance: convergence identifies robust, recurrent sites, while divergence exposes algorithmic sensitivity to transient or cryptic features of the conformational landscape. Early consensus-based approaches [59] demonstrated that combining predictions from multiple algorithms can improve binding site detection accuracy in a static context by leveraging complementary strengths across methods.

Here, we extended this logic to dynamic conformational ensembles, presenting a framework that combines multi-tool spatial consensus with temporal persistence analysis, and validating the approach across two structurally and functionally distinct systems. We first evaluated the concordance between different tool categories and, interestingly, we observed that the consensus was not merely a quality filter, but also an informative descriptor of pocket behavior, helping to distinguish recurrent binding regions from more transient and conformation-dependent sites. Taken together, these results do not point to a single superior tool but rather underscore the complementary nature of the selected approaches: each algorithmic family captures a distinct side of the binding landscape, and it is precisely their divergence that makes their combined use informative. Specifically, inter-tool correlation analysis performed on the GLUT1 ensemble revealed a variable but interpretable landscape of methodological agreement. AutoSite achieved the highest pairwise values across all comparisons, likely due to the spatially diffuse nature of affinity grid predictions, while moderate agreement was observed between SiteHound and P2Rank (r = 0.320) and between Fpocket and SiteFerret (r = 0.242). In contrast, the AR ensemble displayed a substantial correlation increase, with DeepPocket reaching r = 0.590 with SiteHound and r = 0.573 with P2Rank. A particularly informative example concerns the relationship between Fpocket and DeepPocket, which is the lowest pairwise correlation involving Fpocket, despite DeepPocket using Fpocket as its upstream detector. This discrepancy reflects the DeepPocket architecture, where Fpocket-derived candidates are rescored by a 3D convolutional neural network and subsequently refined through volumetric segmentation. Consequently, DeepPocket does not simply inherit the original Fpocket pocket definitions but generates a learned three-dimensional representation of the binding site. This transformation produces a qualitatively different description of the binding landscape, reducing centroid-based correlation with its upstream detector while highlighting complementary spatial features.

Our spatial consensus analysis, based on centroid-to-centroid matching of density -based clusters, was implemented to complement the correlation framework by directly evaluating whether different algorithms converged on the same physical binding regions. Within methodological categories, the energetic category (AutoSite-SiteHound) showed the highest number of matched clusters in both phases (9 in phase 1, 13 in phase 2), while the ML/DL category (P2Rank-DeepPocket) showed the lowest overlap (3 and 4 clusters respectively), consistent with the structural differences between a surface-point Random Forest classifier and a volumetric U-Net segmentation, which produced only partial spatial agreement. Furthermore, the AR dataset provided an informative test case for cryptic pocket sensitivity, with energetic category identifying a consensus pocket sharing four residues with the known Leu300-driven specificity pocket, demonstrating that probe-based interaction mapping can anticipate cavity formation before a geometric concavity is fully established. As observed in GLUT1, the ML/DL category achieved only partial overlap, while the geometric category detected none, reproducing the same energy > ML/DL > geometry sensitivity gradient across both systems.

The inter-category consensus analysis, requiring agreement across at least four out of six tools spanning distinct methodological classes, identified the most robust and biologically meaningful binding regions across the conformational ensembles. In GLUT1, nine high-confidence clusters were found in phase 1, the majority supported by five tools, with two clusters (*clusters 22* and *2*3) representing the only sites that included DeepPocket predictions and *cluster 23* being the sole cluster in phase 1 spanning all three methodological categories simultaneously. The same criteria yielded 11 clusters in phase 2, with *cluster 36* standing out as the only site in either phase where all six tools agreed simultaneously. From a structural perspective, the consensus pockets revealed a clear reorganization of the GLUT1 binding landscape during transport. While phase 1 was dominated by consensus pockets located within the transmembrane core of the protein (*clusters 29, 23, 14,* and *1*), phase 2 revealed an additional consensus pocket in the upper transmembrane domain (*cluster 7*), a finding somewhat unexpected given the inward-open conformation sampled in this phase. Importantly, a central transmembrane pocket remained conserved across both phases (*cluster 36 in phase 2* occupies a central position spatially coincident with *cluster 14* in phase 1), while consensus regions (*clusters 15* and *9* in *phase 1* and *clusters 30* and *16* in *phase 2,* respectively) involving the intracellular helical (ICH) domain suggested the existence of persistently accessible surfaces with potential allosteric relevance. The identification of a consensus pocket (*cluster 12)* containing Asn288, a residue implicated in both cytochalasin B and synthetic inhibitor binding, further linked the consensus framework to experimentally validated pharmacological regions. (*See Fig.S4 Supp. Materials*) This overlap was limited to a single residue because the consensus analysis operated on cluster centroids rather than on full pocket volumes. Overall, the persistence of a central transmembrane site, the continued accessibility of the ICH domain, and the dynamic redistribution of extracellular pockets supported a model in which GLUT1 druggability is not confined to a single static binding region but is redistributed across the protein surface as a function of the conformational state sampled during transport. The AR inter-category analysis further validated the proposed consensus framework, identifying two regions in the area of the known cryptic pocket [60]. *Cluster 7* shared a residue with the target site, while *cluster 4* localized to the correct protein region despite lacking exact residue overlap. This spatial convergence across methodologically diverse tools confirmed that inter-category consensus effectively narrows the search space to biologically relevant regions even in the absence of a fully open cryptic cavity.

A key contribution of our framework is the introduction of temporal persistence as a dynamic complement to spatial consensus, enabling the distinction between constitutive and transient binding sites. Combining Strict persistence, based on continuous frame-by-frame detection, and Soft persistence, allowing short detection gaps, proved essential to capture the full spectrum of pocket dynamics observed across the GLUT1 trajectory. In phase 1, uniformly high soft persistence values (80–100%) with low inter-tool variability confirmed that the outward-open to outward-occluded transition is dominated by stable, recurrently accessible binding regions. Phase 2 presented a markedly different profile, with a wide range of soft persistence values (22–94%) and a pronounced dissociation between strict and soft modes identifying several sites as transient. The observation that these sites were consistently detected by at least four independent tools reduces the likelihood that they arose from algorithm-specific artifacts. Together, inter-tool consensus and non-zero soft persistence provided a practical operational framework for prioritizing potentially relevant transient binding sites. This distinction has direct implications for drug design, with high-persistence sites, such as *cluster 29* in phase 1 and *cluster 36* in phase 2, that represent primary candidates for downstream structure-based studies, while transient pockets may require ligands capable of stabilizing the conformational states in which they become accessible, as demonstrated by allosteric modulator campaigns targeting cryptic sites in otherwise undruggable proteins. Furthermore, in AR dataset both consensus pockets identified near the Leu300 specificity pocket achieved high soft persistence (71–74%) despite the discontinuous nature of cryptic site opening, supporting the Soft persistence criterion as a sensitive detector of biologically meaningful transient binding events.

The temporal dimension captured by persistence analysis was further corroborated by the volumetric consensus strategy. The comparison between HDBSCAN clustering and IoU analysis revealed not only spatial agreement but also complementary sensitivities to different aspects of pocket dynamics. Several sites in phase 2 showed high HDBSCAN soft persistence alongside low IoU occupancy, revealing a fundamental difference in what the two methods measure: HDBSCAN integrates recurrent but discontinuous detection signals across the trajectory, making it more sensitive to transiently sampled pockets, while IoU captures volumetric overlap within discrete trajectory windows and therefore underrepresents sites that are visited frequently but briefly. Rather than reflecting a methodological inconsistency, this divergence highlights the complementary nature of the two approaches and argues for their joint use as a more complete descriptor of binding site dynamics than either method alone.

### Limitations

Despite the robustness of the proposed framework, several limitations must be acknowledged. First, the framework evaluated agreement among pocket detection algorithms rather than the absolute correctness of individual predictions. Unlike benchmarking studies based on known protein cavities, dynamic conformational ensembles generally lack a definitive ground truth for transient pocket occurrence. Consequently, inter-tool consensus should be interpreted as an indicator of methodological convergence and robustness. Second, the persistence thresholds adopted here, 20% for strict and 10% for soft mode, were derived from MDpocket-based criteria and represented a pragmatic choice rather than a universally validated cutoff. Different proteins and simulation timescales may require adjusted thresholds, and the optimal values are likely system-dependent. Third, while the dual consensus framework reduces false positives by requiring inter-tool agreement, it may simultaneously suppress genuine low-confidence sites that are detected by only one or two tools. In highly dynamic or intrinsically disordered regions, biologically relevant pockets may systematically fall below the four-tool agreement threshold, leading to their exclusion from the final set. Finally, the centroid-based spatial matching criterion, while practical and computationally efficient, does not fully account for pocket shape or volume. Two clusters within the 6.5 Å threshold may share a centroid position while representing geometrically distinct cavities, a limitation partially addressed by the independent IoU analysis but not fully resolved.

## CONCLUSIONS

The systematic identification and prioritization of binding sites in dynamic protein systems still represent a fundamental challenge in computational drug discovery. Here, we reviewed and tested multiple pocket detection algorithms, highlighting how pocket detection in dynamic protein systems depends not only on the conformational ensemble being explored, but also on the methodological perspective used to interrogate it. Geometric, energetic, and ML/DL-based approaches frequently identify overlapping yet non-identical binding regions, reflecting the different assumptions underlying each algorithm. Consequently, no single tool appears sufficient to fully capture the complexity of dynamic binding landscapes.

More broadly, this framework aligns with our wider bioinformatics effort to convert complex biological information into actionable translational hypotheses, from pathway modeling and scalable computation [61,62] to regulatory annotation [63], RNA-mediated target prioritization [64], molecular biomarker discovery [65–69], and genotype–phenotype interpretation in rare or genetically defined diseases [70–76]. In this context, dynamic pocket detection adds a complementary structural layer, helping to connect disease-associated proteins and variants with potential druggable sites for future therapeutic exploration.

By applying representative methods from multiple algorithmic classes to two structurally distinct systems, we show that combining spatial consensus with temporal persistence analysis provides a practical framework for prioritizing recurrent and transient binding regions while reducing method-specific biases. Complementary to MD-focused approaches such as MDpocket, which quantify pocket occupancy along a trajectory, this strategy emphasizes methodological convergence as an additional source of information for the interpretation of dynamic pockets.

In conclusion, this work establishes a transferable and extensible computational pipeline that does not require prior knowledge of binding site location, making it applicable to the growing class of proteins previously considered undruggable. As enhanced sampling methods continue to expand the accessible conformational space of challenging targets, and as structural databases increasingly incorporate dynamic information, frameworks capable of extracting druggability signals from conformational ensembles will become central tools in early-stage drug discovery. We anticipate that the integration of shape-based descriptors, adaptive consensus criteria, and machine learning-based threshold calibration- outlined here as future directions - will further extend the sensitivity and generalizability of this approach, ultimately bridging the gap between protein dynamics and actionable pharmacological targets.

## Supporting information

command lines

