## Supplementary Tables for "Dynamic consensus pocket detection across molecular dynamics ensembles reveals persistent and transient druggable sites"

**TABLE 1.** Values of the min_cluster_size parameter used in the HDBSCAN clustering algorithm for binding site prediction.

| **TOOLS** | **PHASE 1** | **PHASE 2** |
| --- | --- | --- |
| AutoSite | 40 | 100 |
| Fpocket | 40 | 125 |
| P2Rank | 40 | 120 |
| SiteHound/EasyMIFs | 40 | 60 |
| SiteFerret | 20 | 75 |
| DeepPocket | 30 | 100 |

**TABLE 2**. Number of HDBSCAN clusters identified by each tool on the AR conformational ensemble (59 frames).

| **Tool** | **Methodological Category** | **N. Clusters** |
| --- | --- | --- |
| AutoSite | Energy-based | 13 |
| P2Rank | Machine Learning | 5 |
| DeepPocket | Deep Learning | 8 |
| Fpocket | Geometry-based | 17 |
| SiteFerret | Geometry-based | 7 |
| SiteHound | Energy-based | 48 |
