## Supplementary material for "Dynamic consensus pocket detection across molecular dynamics ensembles reveals persistent and transient druggable sites": command lines

**P2RANK**

- prank predict -f <pdb_file> - o <output_dir>

**SITEFERRET**

- python3 -m siteferret <pqr_file>

**FPOCKET**

- fpocket -f <pqr_file>

**AUTOSITE**

- PYSH_1 = "/home/ADFRsuite_x86_64Linux_1.0/bin/pythonsh”

PREP_REC = “/home/ADFRsuite_x86_64Linux_1.0/CCSBpckgs/AutoDockTools/Utilities24/prepare_receptor4.py”

ASPY = “/home/ADFRsuite_x86_64Linux_1.0/CCSBpckgs/AutoSite/bin/AS.py”

cmd_prep = f"{PYSH_1} {PREP_REC} -r {pdb_file} -A none -o {pdb_file}qt"
cmd_as = f"{PYSH_1} {ASPY} -r {pdb_file}qt -o {folder_name}/"

**EASYMIFS**

- python /home/easymifs-master/Easymifs.py <pqr_file> <probe>

**SITEHOUND**

- sitehound -f = <pdb_file_CMET.dx> - t = easymifs -l = average -e = - 8.9 -s = 7.8

**DEEPPOCKET**

- python /home/DeepPocket/predict.py -c /home/DeepPocket/DeepPocket/classification_models/first_model_fold6_best_test_auc_196001.pth.tar	-s /home/DeepPocket/DeepPocket/segmentation_models/seg0_best_test_IOU_91.pth.tar -p protein_<frame_num>.pdb -r 15	-t 0.01 --mask_dist 5.0
