## Supplementary Information for "Dynamic consensus pocket detection across molecular dynamics ensembles reveals persistent and transient druggable sites"

Scheda 1

Definition of the Spatial Matching Criterion

Given the high dimensionality and inherence noise of this raw dataset, a spatial correspondence criterion was applied to determine when pocket centroids predicted by different tools were considered to represent the same binding region. Three centroid distance thresholds were systematically evaluated (4, 7, and 10 Å) to span the range from near-perfect overlap to plausible spatial proximity **(Figure S1)**.

The number of matches increased sharply between 4 and 7 Å, suggesting that this range effectively captures high-confidence, near-overlapping predictions.

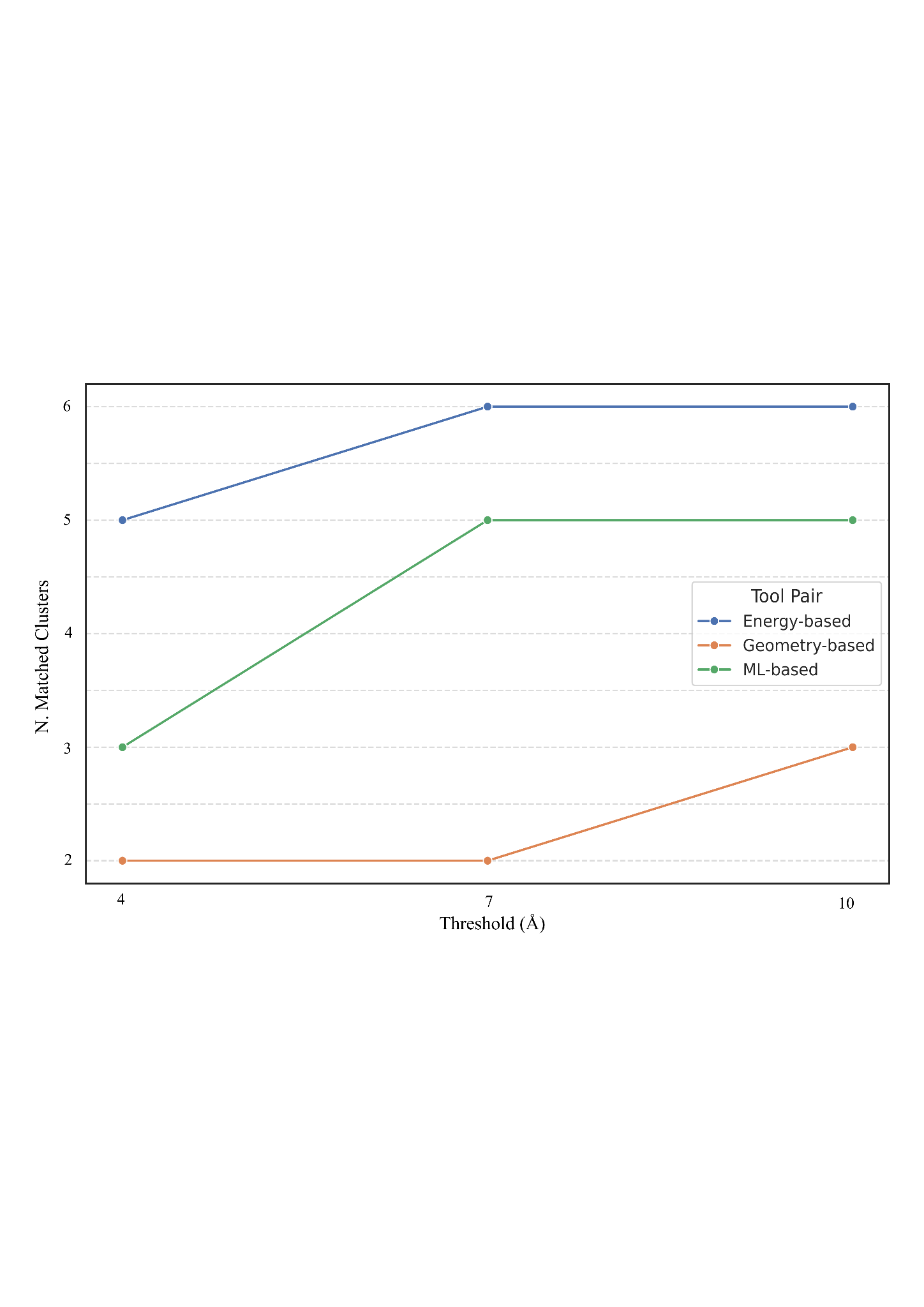

**Figure S1** Shows the variation in the number of clusters identified in both the intra-category consensus analyses during Phase 1 of the molecular dynamics trajectory.

In contrast, the increase from 7 to 10 Å was minimal across all tool pairs. Moreover, matches appearing only at 10 Å were often spatially distant and inconsistent with genuine pocket overlap upon three-dimensional structural inspection (**Figure 1**), suggesting they may represent spurious matches between spatially distinct binding regions.

Based on these findings, a distance cutoff of 6.5 Å was selected as the primary spatial matching criterion distance cutoff. This value was selected as a conservative midpoint within the 4–7 Å range, balancing sensitivity to genuine spatial overlap with robustness against false positive matches.

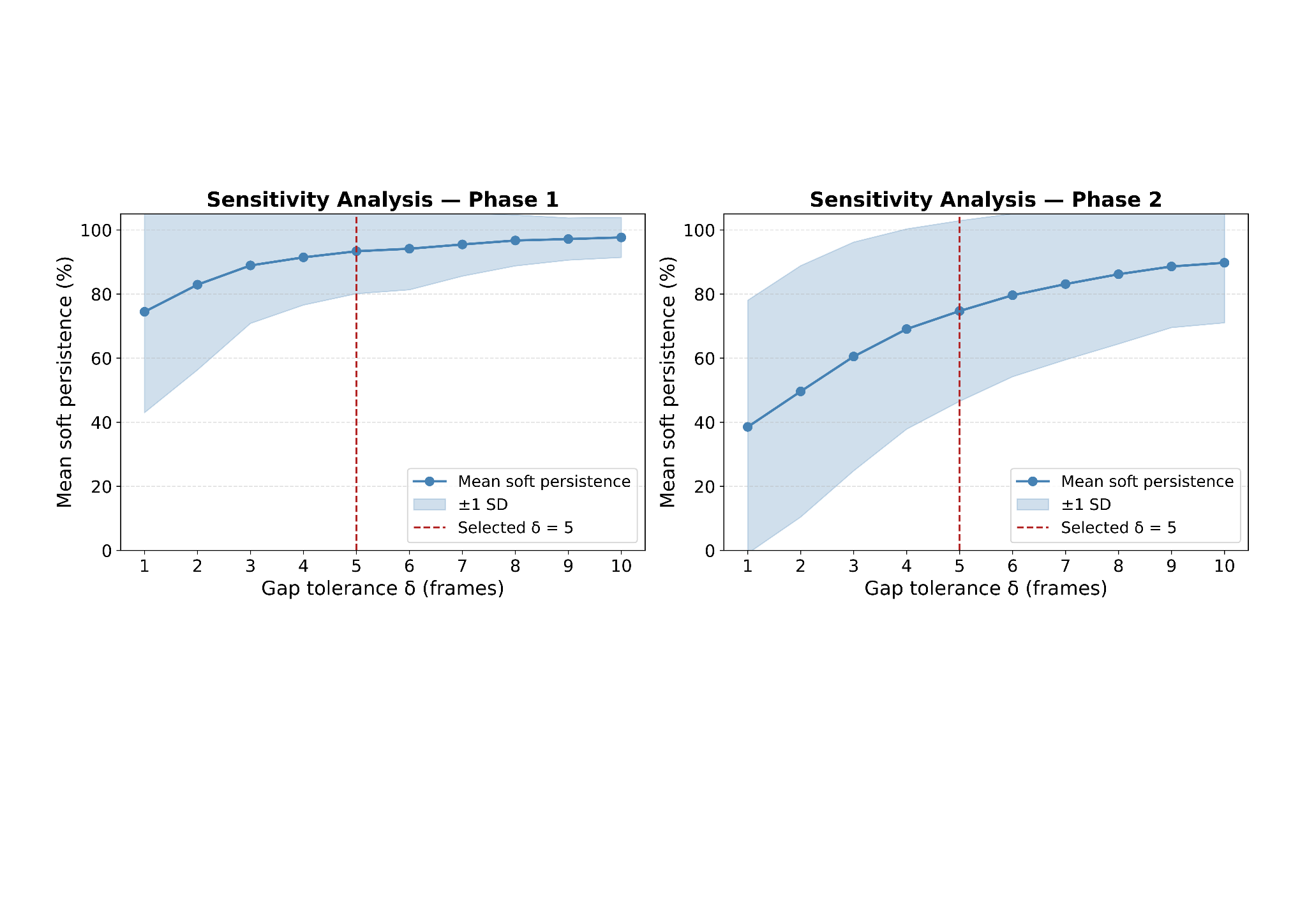

**Figure S2.** Sensitivity analysis of the gap tolerance parameter δ on HDBSCAN-based soft persistence. Mean soft persistence (blue line) computed across all consensus cluster–tool assignments is shown as a function of the gap tolerance parameter δ (1–10 frames), for the GLUT1 conformational ensemble in Phase 1 (left) and Phase 2 (right). The shaded area indicates ±1 standard deviation across consensus cluster–tool pairs. The red dashed line marks the value δ = 5 adopted in the main analysis. In both phases, persistence values rise sharply for δ < 5 and reach a plateau beyond this threshold, with only marginal gains observed at larger δ. This pattern indicates that δ = 5 is the minimum value sufficient to absorb brief detection discontinuities without artificially inflating persistence estimates.

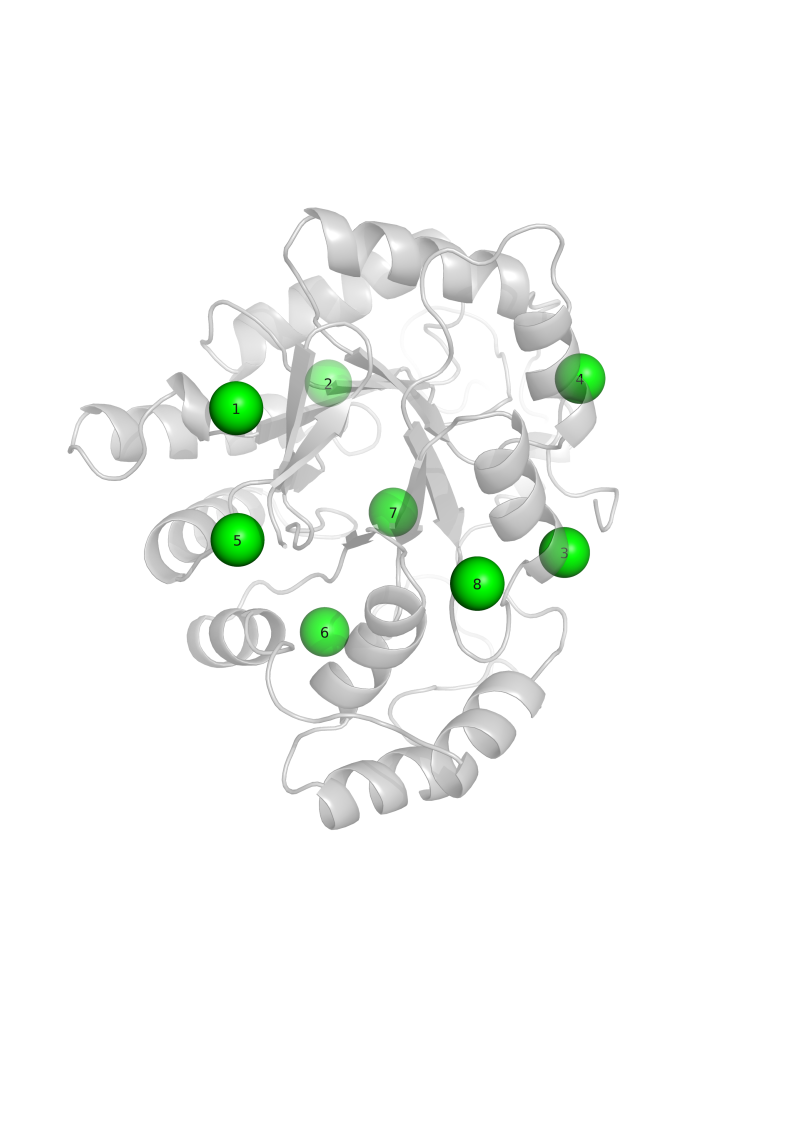

### **Figure S3.** Three-dimensional localization of HDBSCAN consensus binding sites on the Androgen Receptor. Green spheres indicate the centroid positions of the eight consensus clusters mapped onto the AR structure (cartoon representation, gray).

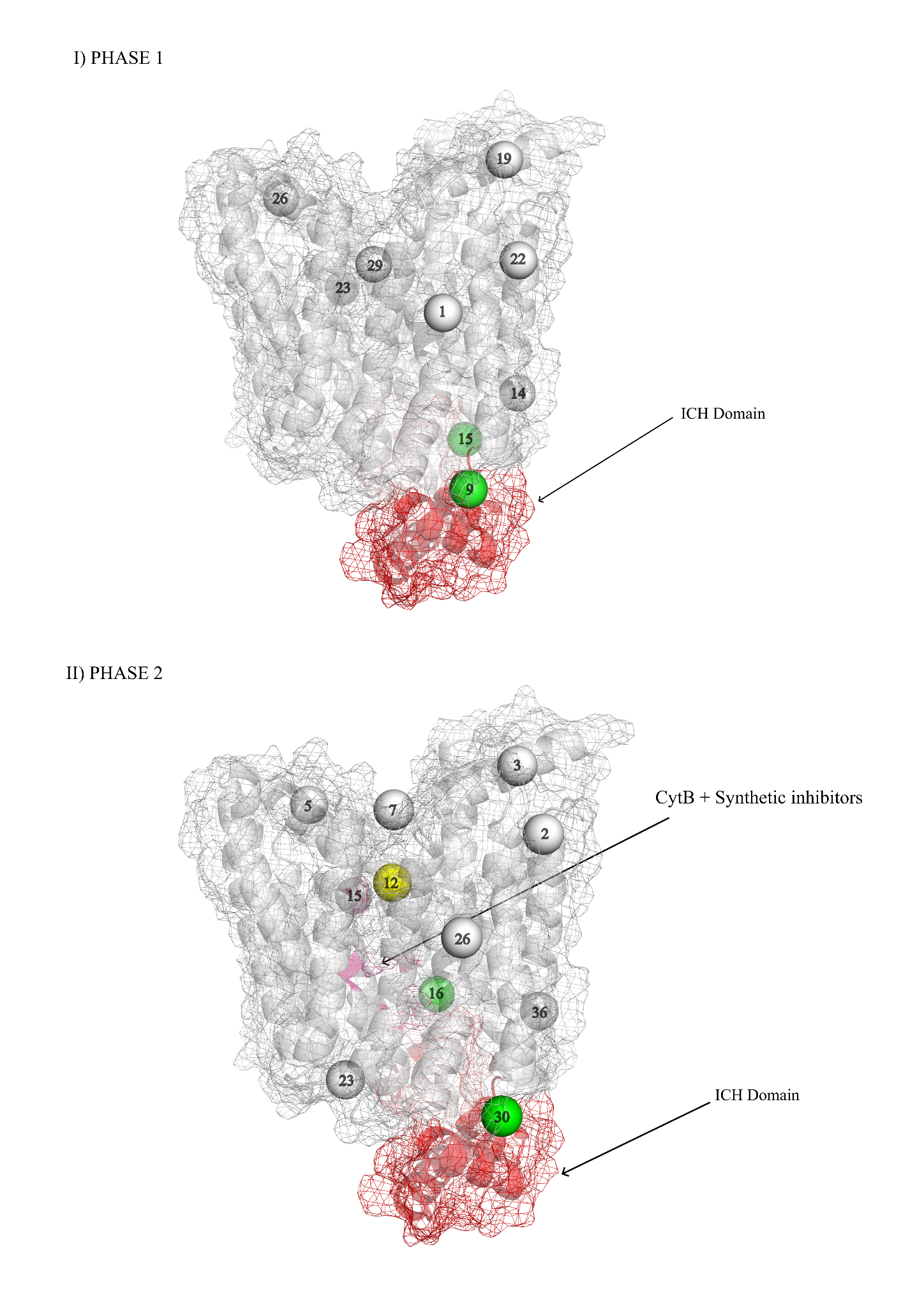

**Figure S4.** Inter-category consensus pockets mapped onto the GLUT1 molecular surface. Pocket centroids are shown as spheres: grey, no known functional overlap; green, pockets within the ICH domain; yellow, pocket overlapping with the cytochalasin B and synthetic inhibitor binding site. The ICH domain is highlighted as a red mesh. I) Phase 1**:** Cluster 9 and Cluster 15 (green) localize within the ICH region on the cytoplasmic face. II) Phase 2: Cluster 30 and Cluster 16 (green) recapitulate the ICH localization; Cluster 12 (yellow) maps onto the pharmacologically validated region containing Asn288.
